# Brain Network Mechanisms of General Intelligence

**DOI:** 10.1101/657205

**Authors:** Chandra Sripada, Mike Angstadt, Saige Rutherford, Aman Taxali

**Affiliations:** Department of Psychiatry, University of Michigan, Ann Arbor, MI

**Keywords:** general intelligence, fronto-parietal network, default network, Human Connectome Project, Adolescent Brain Cognitive Development (ABCD) Study, N-back task, resting state fMRI

## Abstract

We identify novel mechanisms of general intelligence involving activation patterns of large-scale brain networks. During hard, cognitively demanding tasks, the fronto-parietal network differentially activates relative to the default mode network, creating greater “separation” between the networks, while during easy tasks, network separation is reduced. In 920 adults in the Human Connectome Project dataset, we demonstrate that these network separation patterns across hard and easy task conditions are strongly associated with general intelligence, accounting for 21% of the variance in intelligence scores across individuals. Moreover, we identify the presence of a crossover relationship in which FPN-DMN separation profiles that strongly predict higher intelligence in hard task conditions reverse direction and strongly predict lower intelligence in easy conditions, helping to resolve conflicting findings in the literature. We further clarify key properties of FPN-DMN separation: It is a mediator, and not just a marker, of general intelligence, and FPN-DMN separation profiles during the task state can be reliably predicted from connectivity patterns during rest. We demonstrate the robustness of our results by replicating them in a second task and in an independent large sample of youth. Overall, our results establish FPN-DMN separation as a major locus of individual differences in general intelligence, and raise intriguing new questions about how FPN-DMN separation is regulated in different cognitive tasks, across the lifespan, and in health and disease.

## Introduction

There is substantial evidence for an overarching general ability involved in performance across a diverse range of cognitive tasks.^1–5^ This ability, which we here refer to as “general intelligence”^6,7^, is a core element of individual differences in psychological functioning and a key contributor to a number of important academic, occupational, health, and well-being-related outcomes.^8–13^ There has thus been longstanding interest in cognitive neuroscience in identifying the brain mechanisms that produce general intelligence.^14–17^

Previous studies have mainly investigated relatively static, enduring features of the brain that correlate with general intelligence, for example brain size^18^, cortical thickness/gray matter volume^19,20^, and white matter structure^21^ (for reviews, see ^14–17^). A relatively small set of studies examined activation of brain regions during cognitive tasks, and they yielded a mixed picture. Some studies found higher activation in executive regions in subjects with higher intelligence (or better task performance)^22–25^, some found lower activation^26–28^, and others found no relationship^29^.

In the present study, we introduce a novel perspective on brain mechanisms of general intelligence, which in addition sheds light on the mixed findings in previous research. Our approach involves two major shifts from prior work. First, we focus not on neural activation at individual brain regions, but rather activation averaged over distributed large-scale brain networks^30–33^, focusing on mean activation in fronto-parietal network (FPN), associated with executive processing and top-down control^34–38^, and default mode network (DMN), associated with spontaneous cognition^39–42^. These networks are known to exhibit antagonistic relationships during externally-focused cognitively demanding tasks^43,44^, with FPN activating in accordance with cognitive demands^45–47^ and DMN correspondingly deactivating^48–50^, which in turn produces greater “separation” in activation levels between the networks.

Second, we examine network activation profiles not in a single task condition but rather comparatively across easy and hard task conditions. Taking this comparative approach allows us to identify a novel mechanism of general intelligence: generation of greater divergence in network activation profiles across task conditions. In particular, we demonstrate the presence of a crossover relationship: Individuals with higher intelligence exhibit smaller separation between FPN and DMN during easy task conditions and larger separation during hard task conditions. Commensurately, they exhibit greater *change* in network separation between these conditions.

Behavioral and imaging data for our main analysis came from the Human Connectome Project (HCP) 1200 release^51,52^ which has 1206 subjects, of which 920 had usable, high quality data for the main analysis in the present study (age mean 28.6, sd 3.7; female 52.9%). We generated general intelligence scores for each subject by performing bifactor modeling on ten behavioral tasks from the HCP dataset, including seven tasks from the NIH Toolbox and three tasks from the Penn Neurocognitive Battery, and established the model has very good fit to the data (see Methods for details on the factor loadings and fit statistics).

The HCP dataset also includes activation data from a number of tasks performed during neuroimaging scanning, including the N-back task, a widely used probe of working memory. In prior work, working memory tasks were often used to investigate brain regions implicated in general intelligence^22,24,29^, due to extensive evidence that working memory and intelligence are closely related^53–56^. In other lines of research, a number of structural and functional imaging studies have highlighted the involvement of FPN and DMN in general intelligence^14,57,58^. Of particular relevance, we recently showed that cognitive tasks that produce greater separation between FPN and DMN activation are more effective for prediction of intelligence^59^. Building on this body of prior work, we aimed to investigate activation patterns during the N-back task that are associated with individual differences in general intelligence, focusing on the role of FPN/DMN separation.

## Results

### 1. FPN/DMN separation in the N-back task is strongly associated with general intelligence

All participants performed an 9.72 minute N-back task in which they are shown a series of pictures, one every 2.5 seconds. In the harder 2-back condition, they respond when the picture shown on the screen is the same as the one two trials back. In the easier 0-back condition, they respond when the picture is the same as the one shown at the start of the block.

Figure 1 shows mean activation for FPN and DMN in the 0-back and 2-back conditions of the N-back task. The figure highlights that the magnitude of the separation between FPN and DMN (i.e., the difference in activation between the two networks) roughly doubles in the 2-back condition (mean separation = 46.4) compared to the 0-back condition (mean separation = 24.1), a highly statistically significant difference (paired *t*(919) = 65.2, *p* < 1×10^16^).

**Figure 1:**
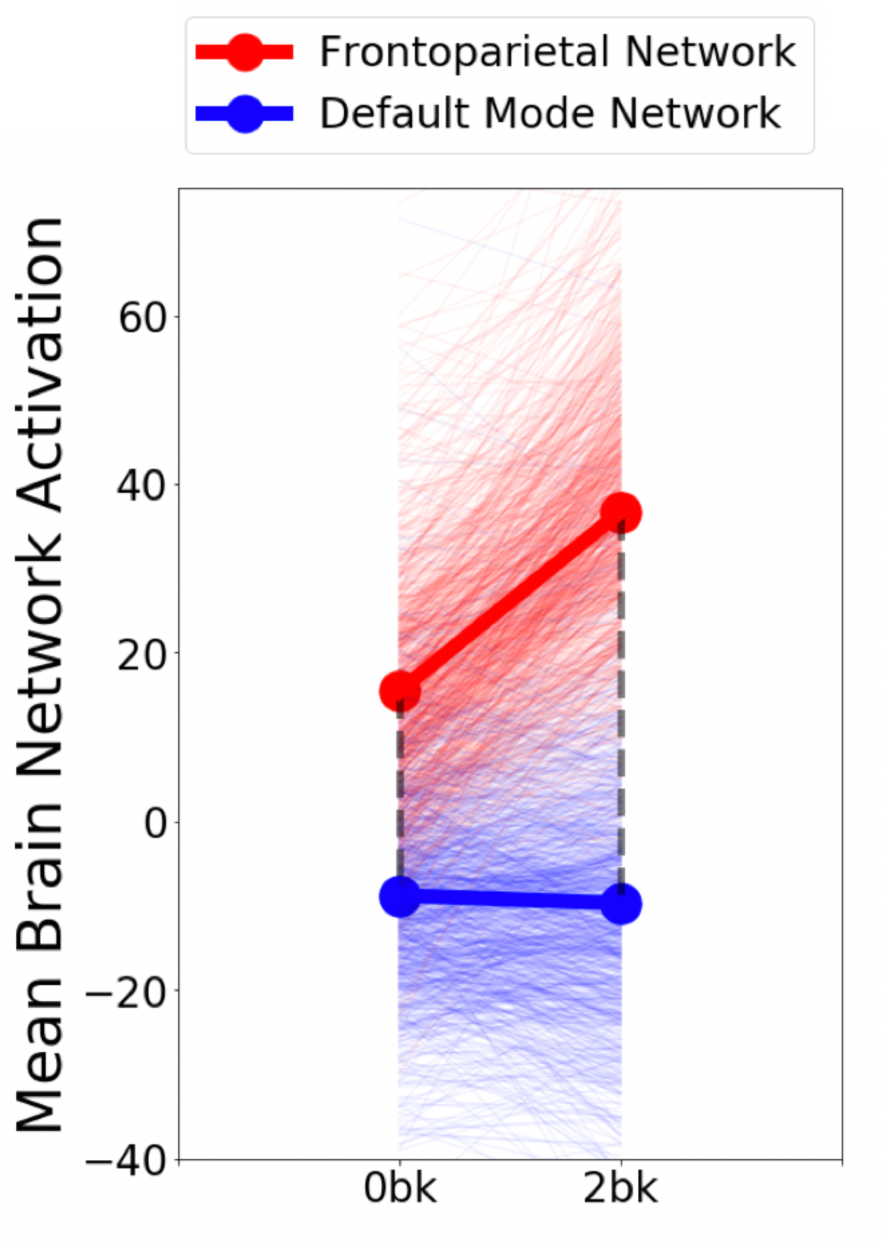
Mean Network Activation for FPN And DMN in the 0-Back And 2-Back Conditions of the N-Back Task. We extracted activation in FPN and DMN during 0-back and 2-back conditions of an N-back task. Separation between FPN and DMN (i.e., the difference in activation between the two networks, shown as dashed lines) roughly doubles from 24.1 in the 0-back to 46.4 in the 2-back. Note: Mean brain network activation is measured in arbitrary units.

To assess the relationship between FPN/DMN separation and individual differences in general intelligence, we constructed a multiple regression model with general intelligence as the outcome and FPN/DMN separation for the 0-back and 2-back conditions as the predictors. Results showed that the overall model was highly statistically significant (F(2,917)=60.5, *p* < 1 × 10−^16^) and explained 20.8% of the variance in general intelligence scores (correlation between model predictions and actual intelligence is 0.46; see Figure 2). Importantly, this effect size is notably large relative to effect sizes typically reported in the “neural correlates of intelligence” field. For comparison, the effect size of brain volume, one of the most studied predictors of intelligence, is typically measured to be roughly a quarter to half this size.^17,18^

**Figure 2:**
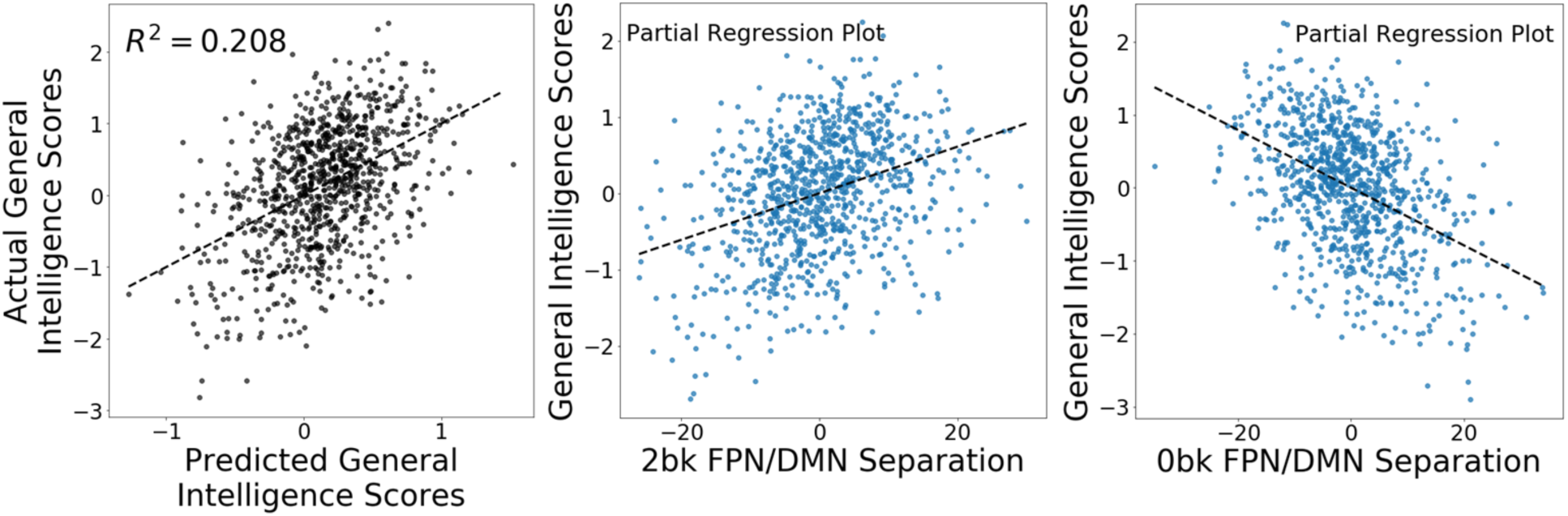
Relationship Between FPN/DMN Separation and General Intelligence in the N-Back Task. (Left Panel) A multiple regression model with FPN/DMN separation values for the 2-back and 0-back conditions as predictors and general intelligence as the outcome explained 20.8% of the variance in general intelligence scores. (**Middle and Right Panels**) The betas in this regression were both highly statistically significant and opposite in sign, indicating that larger FPN-DMN separation predicts higher general intelligence in the 2-back while lower FPN-DMN separation predicts higher general intelligence in the 0-back. Note: Mean brain network activation is measured in arbitrary units.

We additionally examined whether FPN activation alone, rather than FPN/DMN separation, might be an effective predictor of general intelligence. This possibility is motivated by Figure 1, which shows that mean FPN activation increases substantially in the 2-back relative to the 0-back (thick red line), whereas DMN activation remains largely unchanged (thick blue line; note that DMN remains unchanged only when averaging over all the subjects, and there is still substantial variability from subject to subject). We compared an FPN-only model with the FPN/DMN separation model with a likelihood ratio test for non-nested models. Results showed that the FPN-only model was substantially less effective (*z*=6.2, *p* < 10^−8^, r-squared = 8.2%), with r-squared in the prediction of general intelligence decreasing by more than half. This suggests that it is FPN/DMN separation, rather than FPN activation alone, that appears to be most closely linked to general intelligence.

### 2. The relationship between FPN/DMN separation and general intelligence is moderated by task load

In the preceding regression model with general intelligence as the outcome and FPN/DMN separation for the 0-back and 2-back conditions as the predictors, beta weights were highly statistically significant for both predictors (FPN/DMN separation 2-back standardized beta = 0.41; t(917)=11.2, *p* < 1 × 10−^16^; FPN/DMN separation 0-back standardized beta = −0.55; t(917)=15.2, *p* < 1 × 10−^16^). This result indicates the 0-back condition is not simply playing the role of a control condition in this task with respect to intelligence prediction, but rather it contributes independent information of its own. Additionally, the beta weights for FPN/DMN separation in each condition have opposing signs, as shown in Figure 2, middle and right panels, and are significantly different from each other (test of linear contrast: t(917)=14.8, *p* < 1 × 10−^16^). In short then, there is clear evidence of a crossover relationship in which larger FPN-DMN separation predicts higher intelligence in the 2-back while lower FPN-DMN separation predicts higher intelligence in the 0-back.

### 3. FPN/DMN separation values are highly correlated across N-back conditions

We found that FPN/DMN separation values were highly correlated across the 2-back and 0-back conditions (*r*=0.584; *p* < 1 × 10^16^). This perhaps reflects a relatively stable property of the individual to exhibit a characteristic FPN/DMN separation value during the N-back task. Despite being highly correlated with each other, FPN/DMN separation values in the 2-back and 0-back conditions exhibit strongly opposing relationships with general intelligence across these two conditions.

Figure 3 helps clarify these somewhat complex relationships. The dashed line reflects the high correlation between FPN/DMN separation in the 0-back condition and 2-back condition. Individuals with higher intelligence disproportionately reside in the top left “quadrant”. However, because of the strong intercorrelation between 0-back and 2-back FPN/DMN separation values, individuals towards the top on the y-axis tend to be towards the right on the x-axis, and individuals towards the left on the x-axis tend to towards the bottom on the y-axis. Those in the upper left quadrant thus have, compared to expectations implied by the correlation trend line, relatively lower FPN/DMN separation in the 0-back and relatively higher FPN/DMN separation in the 2-back. This pattern, of course, implies greater change in FPN/DMN separation across the two conditions, which we next directly examine and quantify.

**Figure 3:**
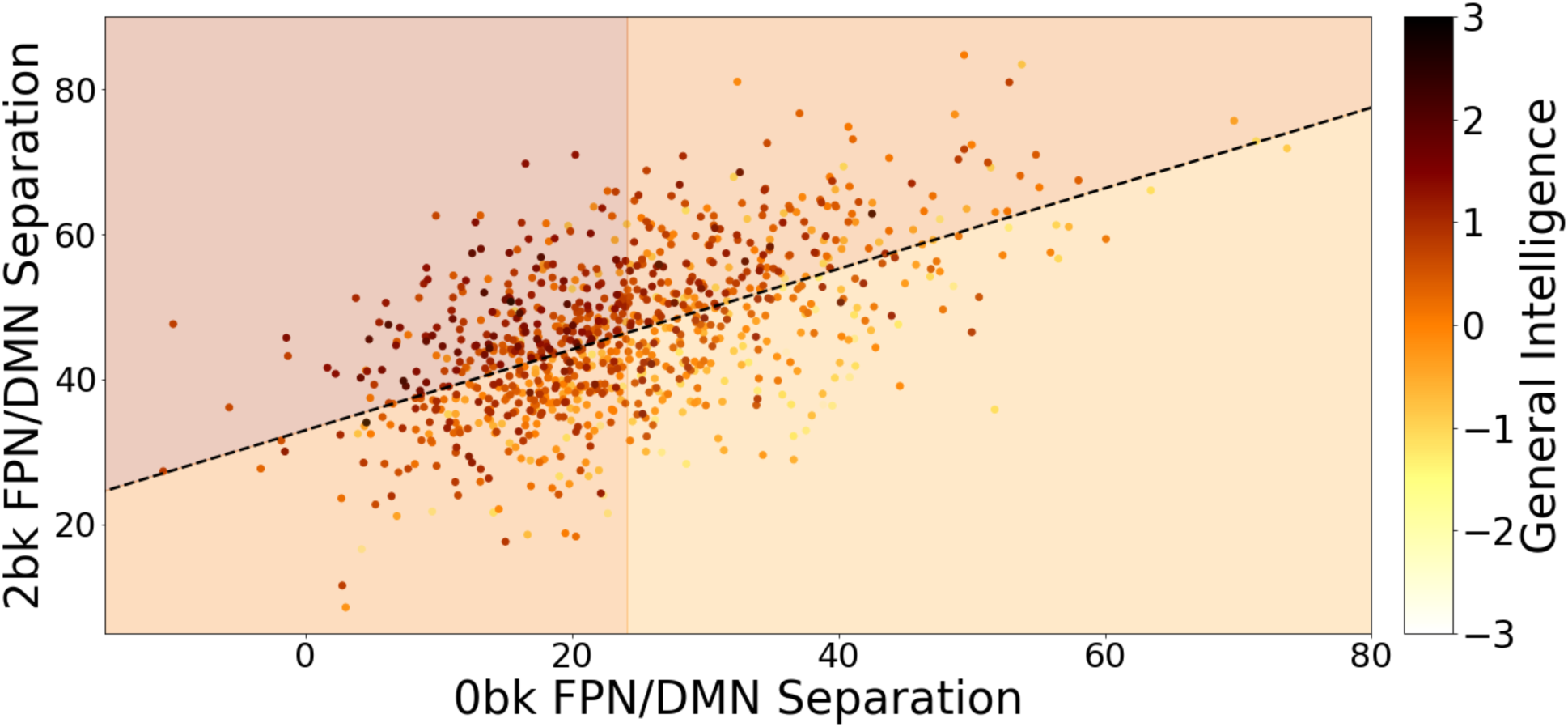
Correlation Between FPN/DMN Separation in the 0-back and 2-back Conditions of the N-Back Task. We observed strong correlation between FPN/DMN separation in the 0-back and 2-back conditions (dashed line). Individuals with higher intelligence (darker red hues) are more concentrated in the upper left quadrant. This quadrant reflects relatively lower FPN/DMN separation in the 0-back and relatively higher FPN/DMN separation in the 2-back compared to expectations implied by the correlation trend line. Note: Shading of quadrants reflects mean general intelligence scores for individuals in that quadrant; FPN/DMN separation is measured in arbitrary units.

### 4. Higher general intelligence is associated with greater change in separation across the 0-back and 2-back conditions

Figure 4 shows differences in FPN/DMN separation across the 0-back and 2-back among participants placed into 10 bins ordered by general intelligence. The figure highlights two interconnected features that characterize those with higher intelligence. First, these individuals tend to have lower FPN/DMN separation in the 0-back condition. Second, these individuals increase their FPN/DMN separation relatively more from the 0-back to 2-back, which is represented in the figure by the lengths of the gray lines. As a result, they have FPN/DMN separation values in the 2-back that are similar to, or slightly higher than, individuals with lower intelligence. Put another way, those with higher general intelligence appear to have a greater ability to change FPN/DMN separation values across task conditions.

**Figure 4:**
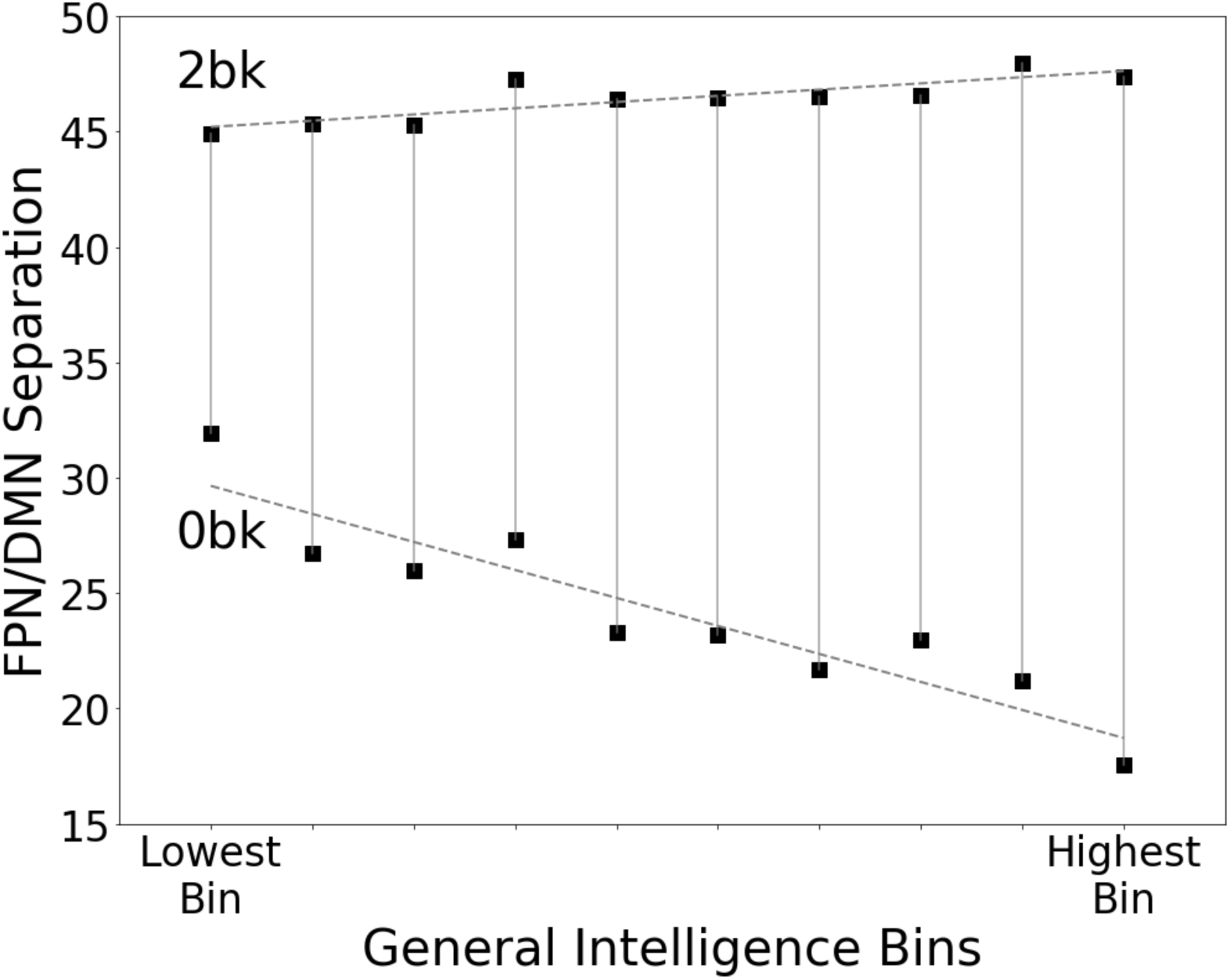
FPN/DMN Separation For 0-back and 2-back by General Intelligence Bin. Participants were placed into 10 bins ordered by general intelligence, and mean FPN/DMN separation values for the 0-back and 2-back conditions for each bin were calculated. The figure shows that those with higher intelligence exhibit two critical properties: (i) They have relatively lower FPN/DMN separation in the 0-back condition; and (ii) They increase their FPN/DMN separation relatively more from the 0-back to 2-back, which is represented in the figure by the lengths of the gray lines. Note: FPN/DMN separation is measured in arbitrary units.

This point can be further illustrated quantitatively. We constructed a multiple regression model in which change in FPN/DMN separation across the 0-back and 2-back was the sole predictor and general intelligence was the outcome. The model’s r-squared was 19.6%, which is very similar to the variance explained by a model with FPN/DMN separation for each condition separately (20.8%). In short then, the change in FPN/DMN separation between the two task conditions, which we hereafter refer to as “delta FPN/DMN separation”, serves as an efficient single-number summary statistic for predicting general intelligence.

### 5. Mediation analysis provides evidence that delta FPN/DMN separation represents a causal mechanism of intelligence

**Figure 5:**
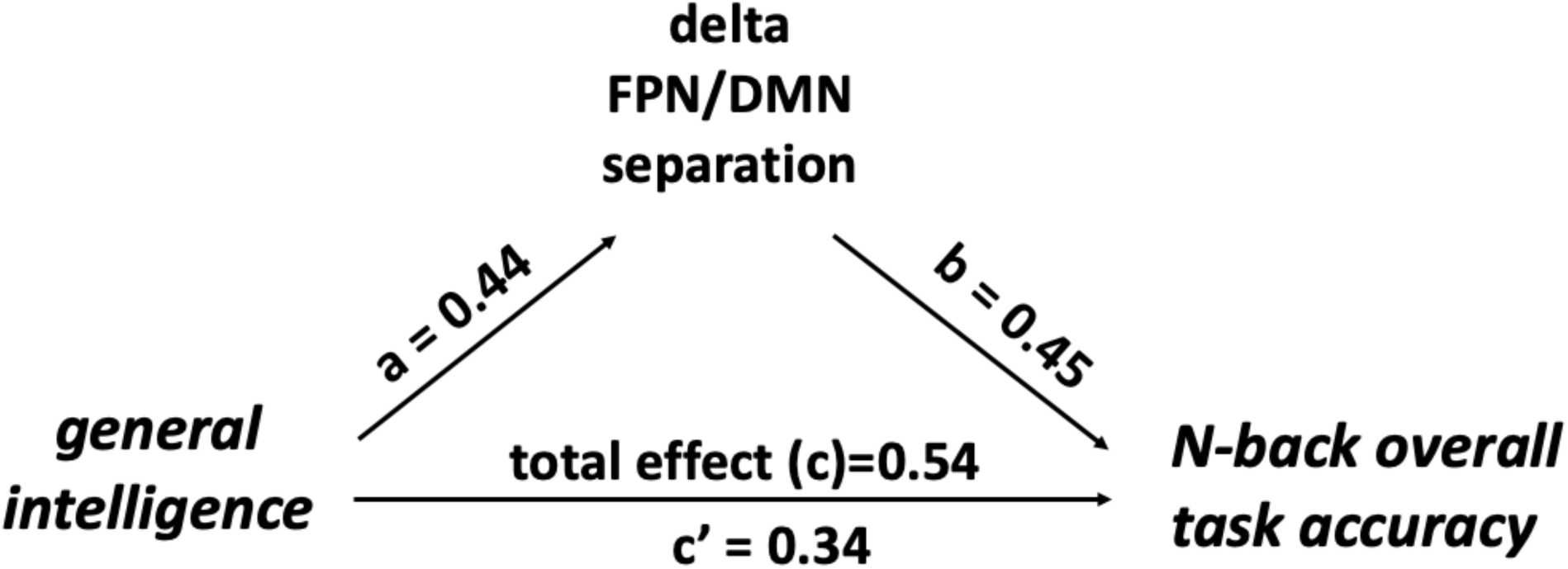
Mediation Model Assessing Causal Role of Delta FPN/DMN Separation in Task Performance. We assessed whether change in FPN/DMN separation from the 0-back to 2-back task (“delta FPN/DMN separation”) serves only as a marker of general intelligence, or whether it mediates the relationship between intelligence and task performance. Results from the mediation analysis showed that 37% of the total effect of general intelligence on N-back task performance is mediated by delta FPN/DMN separation.

We next examined whether FPN/DMN separation is only a marker of general intelligence, or whether it plays a mediating role in producing task performance. We constructed a mediation model with general intelligence as the predictor and N-back overall task performance as the outcome variable. We placed delta FPN/DMN separation as a mediator of this relationship, and assessed mediation with the bootstrapping-based method in the *mediation* package in *R*^60^. Results showed that the mediation pathway was highly statistically significant (*p* < 1 × 10^−8^), and accounts for 37% (95% CI: lower 31%, upper 44%) of the relationship between general intelligence and task performance.

Of note, previous neuroimaging studies^22,29^ examining similar mediation relationships faced a key challenge: they used whole-brain search to identify activation correlates of general intelligence, making mediation statistics involving the regions identified harder to interpret. Our results thus present unusually clear evidence that brain activation patterns mediate the relationship between general intelligence and task performance.

### 6. Delta FPN/DMN separation exhibits high test-retest reliability

A hallmark of general intelligence is that it is exhibits high temporally stability across testing sessions.^61^ In the HCP dataset specifically, using all 46 subjects with available retest data, we found general intelligence scores have a test-retest reliability of 0.78 (ICC type (3,1) in the scheme of Shrout and Fleiss^62^). If FPN/DMN separation is a core mechanism of general intelligence, it too should have similar temporal stability. This is indeed what we observed. In 33 subjects with usable retest data for the N-back task, we found delta FPN-DMN separation has a test-retest reliability of 0.71 (Figure 6).

**Figure 6:**
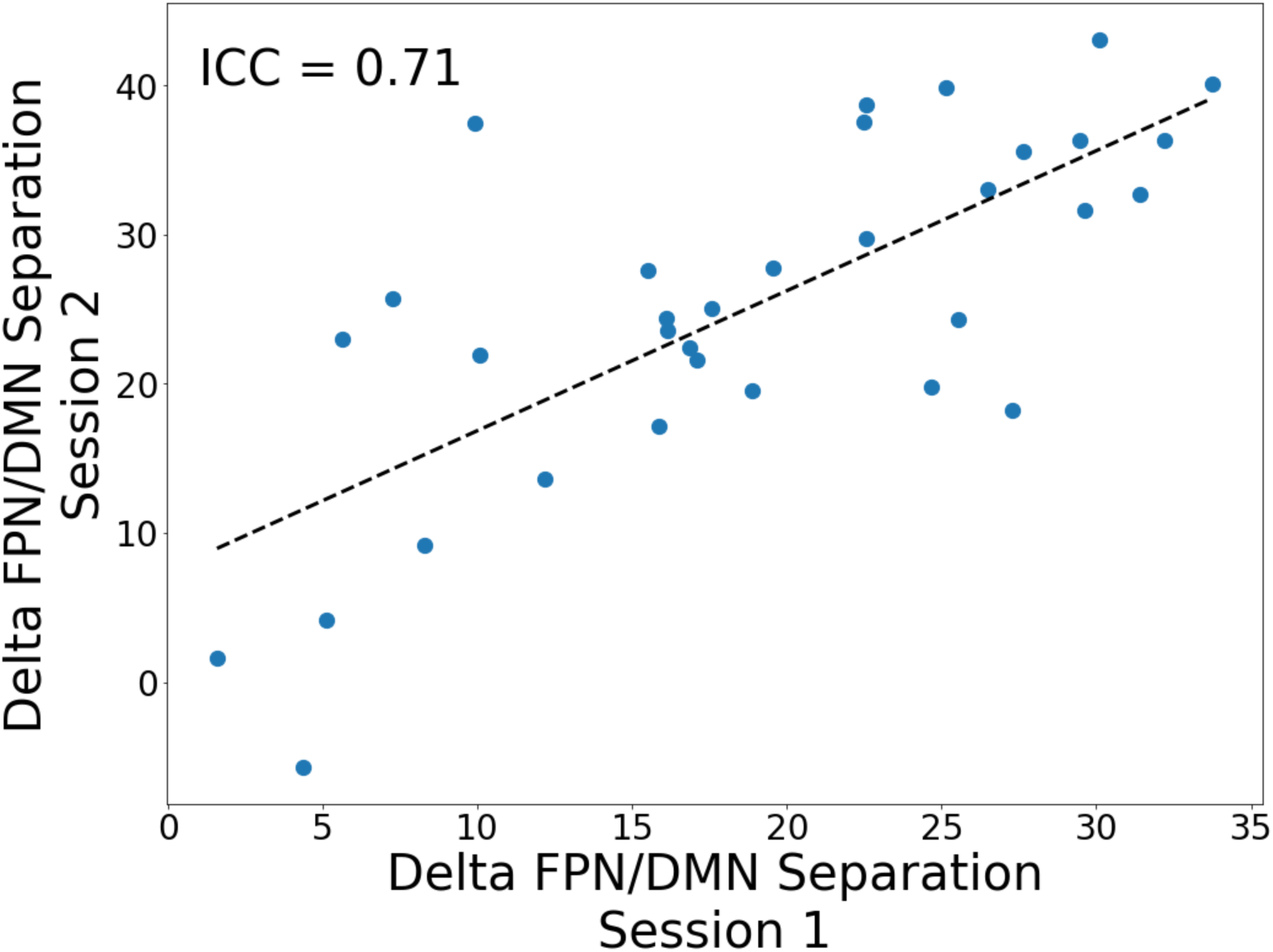
Test-Retest Reliability of Delta FPN/DMN Separation. General intelligence is relatively stable across testing sessions. We found that delta FPN/DMN separation, a proposed brain network mechanism of general intelligence, also exhibits relatively high test-retest reliability. Note: FPN/DMN separation is measured in arbitrary units.

### 7. Functional connectivity patterns involving FPN during the resting state are implicated in individual differences in FPN/DMN separation during the task state

FPN and DMN are large-scale intrinsic connectivity networks that are known to exhibit distinctive patterns of intra- and inter-network functional connectivity during the resting state ^36,63,64^. FPN in particular has widespread connectivity with other networks^65^ and these interconnections are thought to be the source of adaptive control signals that regulate activity in the targeted networks^34,65–68^. We thus hypothesized that individual differences in resting state connectivity patterns, particularly connections involving FPN and DMN, would be predictive of individual-differences in FPN/DMN separation values during the N-back task.

To test this hypothesis, we applied brain basis set (BBS), a validated multivariate predictive modeling approach^59,69,70^ (see Supplement for details) to resting state connectomes from HCP subjects to predict their delta FPN/DMN separation during the N-back task. We identified 834 HCP subjects with both usable N-back data and resting state data, and this set was further partitioned into a train dataset with 744 subjects and an independent test dataset of 90 subjects unrelated to the train subjects and to each other.

Results showed that in the independent held out sample, the correlation between predicted delta FPN/DMN separation scores and actual delta FPN/DMN separation scores was 0.40, a highly statistically significant result (t=4.1, *p* < 0.0001). We then visualized the consensus connectome from this predictive model to identify connections within the connectome that account for the model’s success (see Methods for details on how the consensus connectome is created). We found connections involving FPN and DMN were overrepresented: FPN and DMN make up 12.9% percent of connections in the connectome but constitute 36.3% of the suprathreshold connections.

Next, we assessed the importance of specific networks for predicting delta FPN/DMN separation by repeating the BBS analysis using just two networks at a time for all possible pairs of unique networks. Results showed that prediction accuracy actually improved with just two networks for several network pairs (see Figure 7, right panel), all involving FPN, including: FPN-VAN (*r*=0.50), FPN-DAN (*r*=0.47), and FPN-DMN (*r*=0.45; all *p*’s < 0.0001). These results demonstrate a clear link between connectivity patterns during the resting state and FPN/DMN separation profiles during the task state, and they implicate resting state connectivity patterns of FPN in particular as playing a central role.

**Figure 7:**
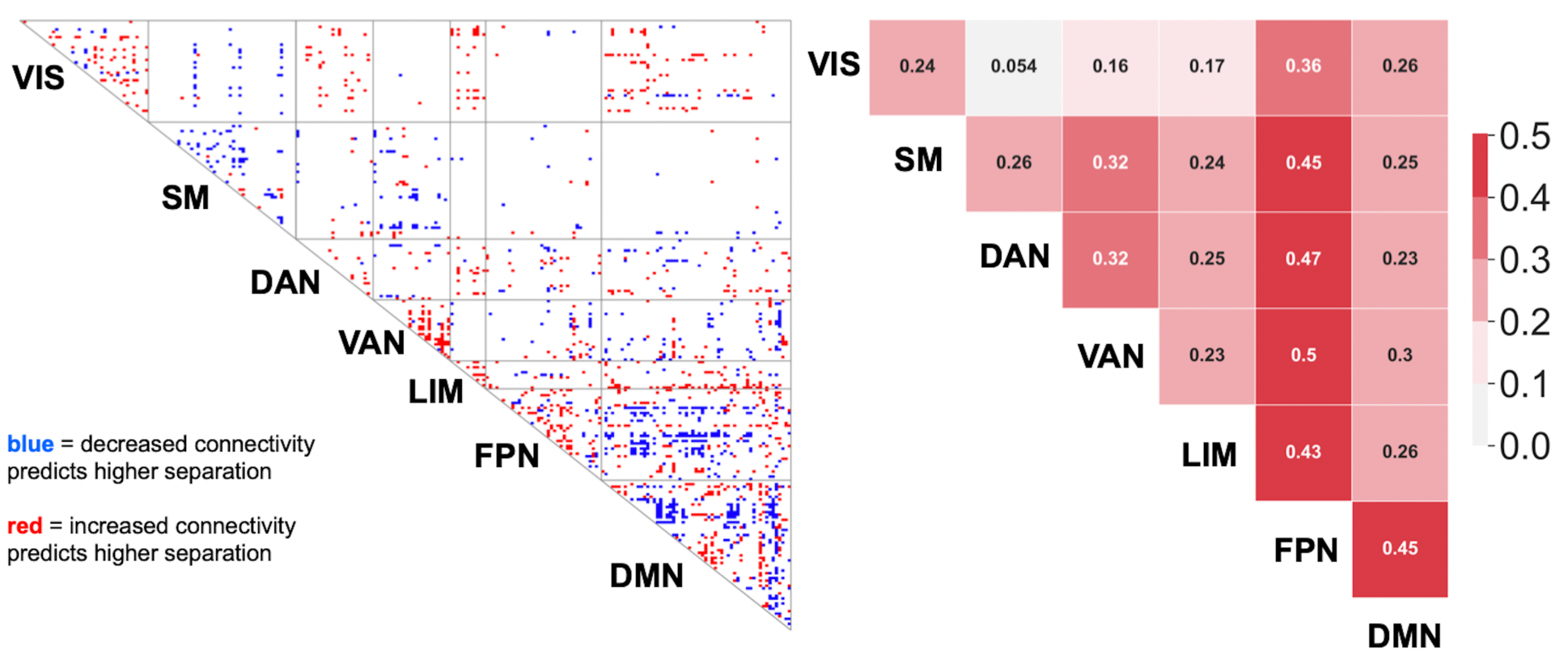
Connections During Resting State That Are Predictive of Delta FPN/DMN Separation During the N-Back Task. Using fully independent train and test samples, we found a predictive model trained on resting state data predicted change in FPN/DMN separation between the 0-back and 2-back conditions (“delta FPN/DMN separation”) with a correlation of 0.40, a highly statistically significant result. (**Left Panel**) Consensus connectome showing connections weighted heavily in the predictive model. (**Right Panel**) Table showing predictive success in follow-up analyses that dropped all networks except two. Several network pairs, all involving FPN, showed strong predictive success, including, FPN-VAN, FPN-DAN, and FPN-DMN.

### 8. Key elements of the preceding analyses replicate in a second HCP task as well as in an independent large youth sample

To assess the robustness of our findings, we repeated the preceding analyses on a second HCP task that involves matched easy and hard conditions, as well as 1,240 subjects in the ABCD youth dataset. In the Relational task (*n*=917; age mean 28.7, sd 3.7; female 52.6%), participants are shown a pair of cue objects. In the “relational” condition, they identify the dimension along which the pair differ (e.g. shape or color), and then determine if a target pair of objects differs along that dimension. In the easier “match” condition, they are given a single cue object and are asked if a member of a target pair of objects matches the cue along a given dimension. Subjects completed 54 total trials over the course of two 2.6 minute runs.

The ABCD sample consists of 9- and 10-year-olds in Release 1.1, which after exclusions has 1,240 usable subjects (age mean 10.1, min 9.0, max 10.9; female 47.5%). All subjects received an extensive neurocognitive battery that included the NIH toolbox measures used in HCP as well as additional measures (e.g., Rey’s learning task, Little Man task). Similar to our approach in the HCP dataset, we fit a bifactor model to these neurocognitive tasks, established there was very good fit with the data, and calculated general intelligence scores for each subject. These subjects also all performed an 9.87-minute N-back task during fMRI scanning, similar to the task used in HCP, and in addition they all received 20-minutes of resting state scanning. Additional details about the ABCD sample demographics, neurocognitive tasks, bifactor modeling and fit statistics, scanning protocols, and methods for constructing imaging maps are described in detail in Methods, as well as in our previous publication.^70^

Figures 8 and 9 and Table 1 show that many of the key results from the HCP N-back dataset were also observed in the two replication datasets, i.e., the HCP Relational task dataset and the ABCD dataset. As shown in Figure 8A and 9A, in both replication datasets we found FPN-DMN separation in the hard and easy conditions were highly predictive of general intelligence scores (Table 1, row 1). In both replication datasets, betas for each predictor were each highly statistically significant and oppositely signed, with higher FPN/DMN separation predicting higher intelligence in the hard condition and lower FPN/DMN separation predicting higher intelligence in the easy condition (Table 1, rows 2 and 3). As shown in Figure 8B and 9B, in both replication datasets, change in FPN/DMN separation across the easy condition and hard condition (indicated by the length of the gray lines) increased with higher intelligence. This is confirmed quantitatively: In both replication datasets, a regression model with general intelligence predicted by delta FPN/DMN separation had an r-squared similar to (or identical to) an analogous model with FPN-DMN separation from each task condition (Table 1, row 4). Additionally, as shown in Figure 8C and 9C, in both replication datasets, delta FPN/DMN separation was found to be highly statistically significant mediator of the relationship between general intelligence and task performance (Table 1, row 5). Finally, in both replication datasets, resting state connectivity patterns were found to be statistically significant predictors of FPN/DMN separation during task (Figures S1 and S2 in the Supplement).

**Figure 8:**
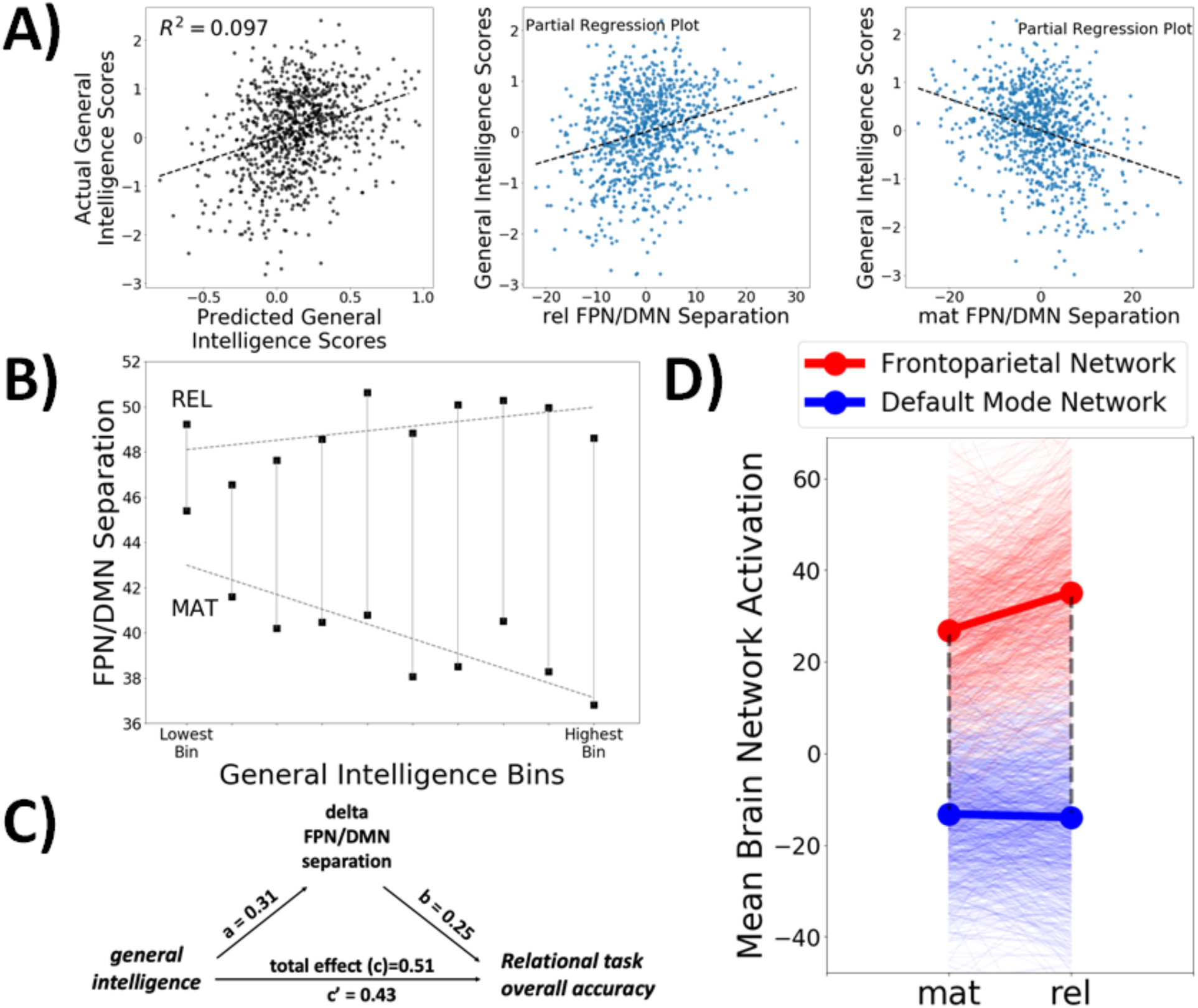
Key Results from the HCP Relational Task. To assess the robustness of our results in a different task, we repeated the preceding analyses on a second HCP task, the Relational task, and we found a highly similar pattern of results. (**A, top panel**) FPN-DMN separation in the relational and match conditions were highly predictive of general intelligence scores. (**A, bottom panel**) There was a clear reversal of the relationship between FPN/DMN separation and general intelligence in the relational versus match conditions. (**B)** Greater change in FPN/DMN separation across the 0-back and 2-back (indicated by the lengths of the gray lines) was associated with higher general intelligence. **(C)** Delta FPN/DMN separation was found to be highly statistically significant mediator of the relationship between general intelligence and Relational task performance, with the mediation pathway accounting for 15% of the total effect. **(D)** Separation between FPN and DMN increases from 40.1 in the match condition to 49.0 in the relational condition, less than half the increase seen in the HCP N-back task. Note: Mean brain network activation and FPN/DMN separation are measured in arbitrary units. mat = match condition; rel = relational condition

**Figure 9:**
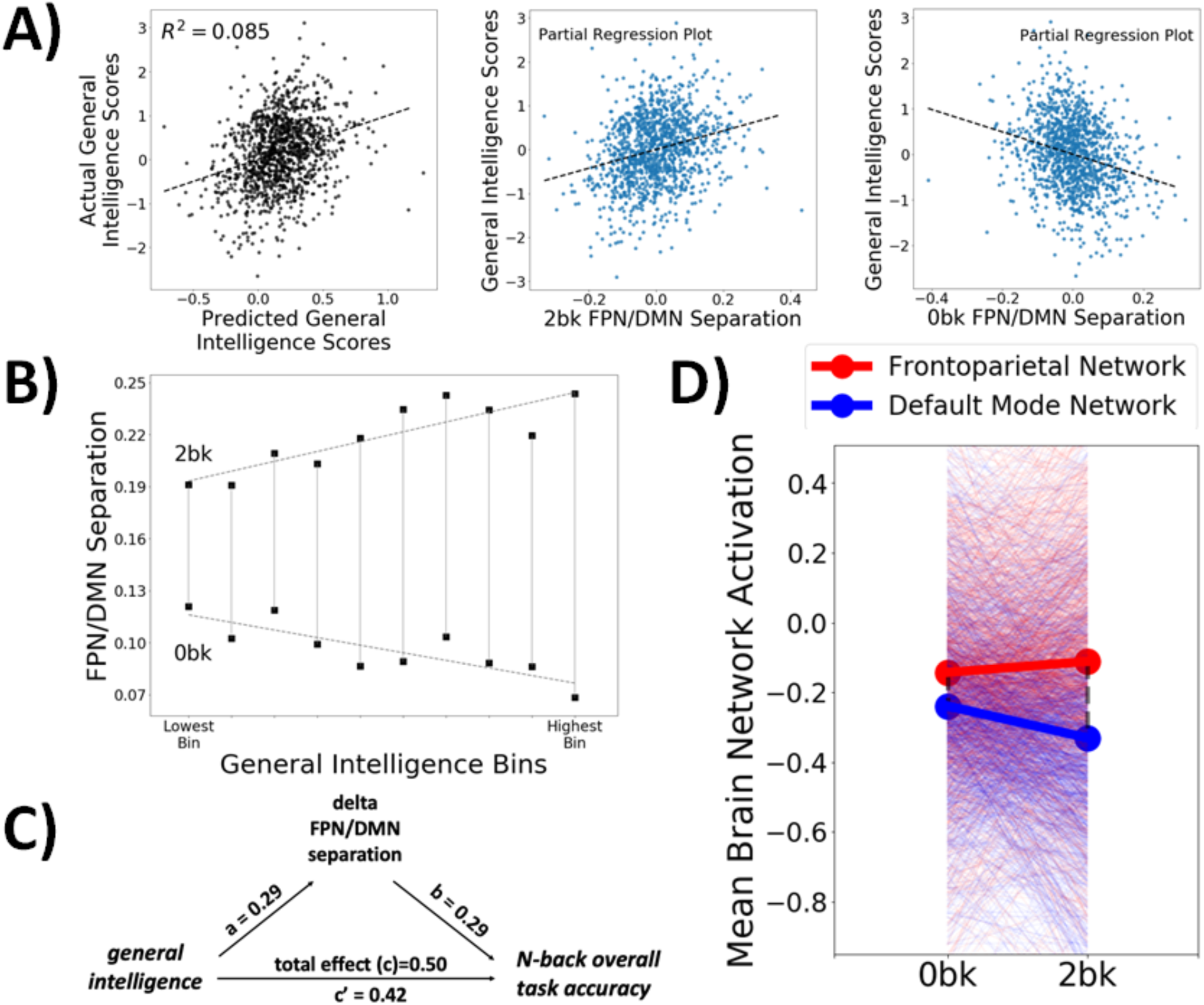
Key Results from 1,240 Youth in the ABCD Dataset. To assess the robustness of our results across different samples, we repeated the preceding analyses performed on the HCP adult dataset in 1,240 9- and 10-year-olds in the ABCD youth dataset, finding a highly similar pattern of results. (**A, top panel**) FPN-DMN separation in the 2-back and 0-back were highly predictive of general intelligence scores. (**A, bottom panel**) There was a clear reversal of the relationship between FPN/DMN separation and general intelligence in the 2-back versus 0-back conditions. (**B) G**reater change in FPN/DMN separation across the 0-back and 2-back (indicated by the lengths of the gray lines) was associated with higher general intelligence. **(C)** Delta FPN/DMN separation was found to be highly statistically significant mediator of the relationship between general intelligence and N-back task performance, with the mediation pathway accounting for 16% of the total effect. **(D)** Separation between FPN and DMN (indicated by the dashed lines) roughly doubles from 0.10 in 0-back to 0.22 in 2-back, similar to change in the HCP N-back task. However, the variability of the increase is much higher in the ABCD youth dataset. Note: Mean brain network activation and FPN/DMN separation are measured in arbitrary units.

**Table 1:** Summary of Key Results from Two Replication Samples. To assess the robustness of our findings, we repeated our analyses on two replication datasets. The first column shows results from the Relational task from the HCP dataset, an abstract reasoning task that also has matched easy and hard conditions. The second column shows results from 1,240 9- and 10-year old participants in the ABCD youth dataset.

| Analysis Element | HCP Relational Task |  |  | ABCD N-back Task |  |  |
| --- | --- | --- | --- | --- | --- | --- |
|  | Effect Size | statistic | *p* | Effect Size | statistic | *p* |
| 1. general intelligence predicted by FPN-DMN separation in hard and easy conditions | R<sup>2</sup>=9.5% | F(2,914)=49.1 | < 1 × 10<sup>-16</sup> | R<sup>2</sup>=8.4% | F(2,1237)=57.2 | < 1 × 10<sup>-16</sup> |
| 2. beta for hard | 0.40 | t(914)=8.4 | < 1 × 10<sup>-16</sup> | 0.31 | t(1237)=9.5 | < 1 × 10<sup>-16</sup> |
| 3. beta for easy | -0.46 | t(914)=-9.8 | < 1 × 10<sup>-16</sup> | -0.30 | t(1237)=-9.3 | < 1 × 10<sup>-16</sup> |
| 4. general intelligence predicted by delta FPN-DMN separation | R<sup>2</sup>=9.5% | F(1,915)=95.7 | < 1 × 10<sup>-16</sup> | R<sup>2</sup>=8.3% | F(1,1238)=10.6 | < 1 × 10<sup>-16</sup> |
| 5. mediation pathway | 15% of total effect | non-parametric | < 2 × 10<sup>-16</sup> | 16% of total effect | non-parametric | < 2 × 10<sup>-16</sup> |

The preceding results demonstrate the generality of the link between FPN/DMN separation and general intelligence, as this link is observed in another quite different type of task (an abstract reasoning task) and in a second, quite different subject population (9- to 10-year olds). But there are also notable differences that bear mentioning. Perhaps the most important is that the effect size of the increase in FPN/DMN separation in the HCP N-back task is substantially larger than in the HCP Relational task as well as the ABCD N-back task, but for different reasons.

In the HCP N-back task, FPN/DMN separation roughly doubles from the 0-back condition to the 2-back condition (Figure 1). In the Relational task, however, FPN/DMN separation increases by only 22% (Figure 8D). This represents a 1.1 standard deviation increase in the Relational task, compared to a 2.1 standard deviation increase in the HCP N-back task. The key difference is that the match condition has an FPN/DMN separation value that is already quite high (Figure 8D), so the overall change in FPN/DMN separation for the Relational task remains quite small. In the ABCD N-back task, on the other hand, FPN/DMN separation roughly doubles (Figure 9D), just as it does in the HCP N-back task. But importantly, due to much greater variability in FPN/DMN separation values in the ABCD dataset, this change between the 0-back and 2-back represents only a 1.2 standard deviation increase in ABCD, i.e., the HCP N-back effect size is nearly twice as large. This difference in effect size is unlikely to be due to differences in tasks: The HCP and ABCD N-back tasks are both block designs with similar stimuli and durations. In the Discussion, we consider other factors that might explain increased FPN/DMN separation variability in the ABCD sample, including subject/scanner heterogeneity in the multi-site ABCD study and immaturity of the brain’s network architecture in 9- and 10-year-olds.

## Discussion

Elucidation of the mechanisms of general intelligence has been a long-standing goal of cognitive neuroscience^14–17^. Our major contribution in this study is to introduce FPN/DMN separation as an important brain network mechanism of general intelligence. We showed this mechanism has a large effect size, accounting for as much as 21% of the variance in general intelligence scores, it is a mediator and not just a marker of general intelligence, and the operation of this mechanism during tasks is at least partly reflected in connectivity patterns of the brain’s intrinsic functional architecture. These results provide a new perspective on how general intelligence is realized in the brain, highlighting the underappreciated role of complex interrelationships between FPN and DMN, two major large-scale brain networks.

The findings from this study should be interpreted in terms of over two decades of observations of antagonistic and cooperative relationships between FPN and DMN across tasks, during neurodevelopment, and in psychiatric disorders. Across a wide range of cognitively demanding tasks, FPN activation increases ^37,38,71,72^, while DMN activation decreases ^73–75^, where the magnitude of FPN activation^45–47^ and DMN deactivation^48–50^ closely tracks the level of cognitive demands. Antagonism between FPN and DMN is also reflected in the phenomenon of lapses of attention, where spikes in DMN activation precede errors^75–78^; during the resting state, where spontaneous slow oscillations in DMN and FPN are anti-correlated^36,44^; and during neurodevelopment, where the two networks become increasingly segregated from childhood to young adulthood^79–81^. Notably, loss of segregation between FPN and DMN is frequently found in mental disorders, including ADHD^81,82^, PTSD^83,84^, and schizophrenia^85,86^. On the other hand, the networks also exhibit cooperative dynamics during certain tasks^87^, for example during contextual recollection^88^, future-directed thought^89,90^ and mind wandering^91,92^. Overall, while complex FPN/DMN interrelationships have long been recognized as a central feature of brain network organization, this study appears to be the first to link the activation profiles of these two networks to general intelligence.

Our results agree with those from previous studies that examined correlations between brain activation patterns during cognitive tasks and general intelligence, but we extend them in a number of important ways, and several specific differences are worth highlighting. First, nearly all previous task-based studies of general intelligence used voxel-wise methods to search across the brain for individual brain regions whose activation correlates with intelligence. Our study encourages a shift in units of analysis from specific voxels or regions to entire brain networks, i.e., distributed large-scale systems that operate as relatively integrated units to perform cognitive functions^93,32^. Second, whereas activation within FPN regions has been repeatedly highlighted in previous studies^22,23,27,28,94^, our study introduces the somewhat novel idea that the critical feature for predicting general intelligence is the joint activation profile of FPN and DMN. Notably, we found a model predicting general intelligence from mean FPN activation alone explained only half the variance of an analogous model using FPN/DMN separation (Results, §1).

Third, we demonstrated robust moderation of brain-intelligence relationships by task difficulty. This finding potentially sheds lights on mixed results observed in previous studies where activation in executive regions was found to correlate positively^22–25^, negatively^26–28^, or exhibit no relationship^29^ with general intelligence. These inconsistent relationships may reflect varying task difficulties across different studies, as well as perhaps varying cognitive abilities of the respective subject populations (e.g., some elite student populations have ceiling performance on tasks that other populations find fairly challenging; see for example ^22,29^). If brain-intelligence relationships are moderated by task difficulty, one would expect to see complex patterns of variation in brain-intelligence relationships across such studies.

Our observation of strong moderation by task difficulty also helps adjudicate between “capacity” and “efficiency” approaches to interpreting task activation correlates of general intelligence^94^. Observations of greater activity in executive regions during cognitive tasks among more intelligent subjects (or subjects with better task performance) led some authors to conclude these subjects have greater executive capacity^22–24^. On the other hand, observations of negative correlations led other authors to favor a neural efficiency explanation^26–28,95^, in which higher intelligence subjects require fewer cognitive resources to achieve similar or better performance. Our results point to both explanations potentially playing a role, with higher intelligence associated with greater capacity in high demand tasks and greater efficiency in low demand tasks^94^. Moreover, they point to a role for a third critical construct: adaptivity. We found those with higher intelligence have better ability to produce larger differences in FPN/DMN separation across the low and high task load conditions, perhaps reflecting an ability to shift between high capacity and high efficiency processing modes.

Our identification of FPN/DMN separation as a key element in individual differences in general intelligence immediately raises a number of new and intriguing questions about the mechanisms that produce individual differences in FPN/DMN separation: How do such differences arise during development?, How are they regulated during task contexts?, Can FPN/DMN separation be modified through cognitive training or pharmacological interventions, among other means?

In the current study, we clarified one key mechanism of individual-differences in FPN/DMN separation: Resting state functional connectivity patterns, especially connectivity of FPN. This result is consistent with emerging models of FPN as a core executive network that flexibly interconnects with various other distributed networks^65^ (including DMN)^87,30^, and which supplies adaptive control signals that modify processing in targeted networks^65–67^. Overall, however, the present study is clearly a starting point rather than a conclusion. By introducing FPN/DMN separation as a new network mechanism strongly linked to general intelligence, the door is opened for future work to elucidate of structural, functional, developmental, and genetic bases that combine to produce adaptive versus maladaptive adjustment in FPN/DMN separation across different cognitive tasks and in health and disease.

We confirmed the robustness of our result by extending them to an additional task in the HCP dataset (one that involves abstract reasoning rather than working memory) as well as an additional sample: 1,240 9- and 10-year-olds in the ABCD dataset. We did observe some diminution of size of the link between FPN/DMN separation and intelligence in these replication datasets. One potential explanation takes note of the fact that the change in FPN/DMN separation observed in these two datasets is substantially smaller—about half the size of the HCP N-back effect size (measured in standard deviation units). It is possible that in tasks that produce less change in FPN/DMN separation across easy and hard conditions, the link between FPN/DMN separation and intelligence is correspondingly muted.

In the N-back task in the ABCD dataset, in particular, reduced change in FPN/DMN separation was due to much greater FPN/DMN separation variability in this dataset, which in turn might be explained in at least two ways. One possibility is that the underlying brain network mechanisms of general intelligence in youth in ABCD are comparable to adults in HCP, but other characteristics of the ABCD sample explain their greater FPN/DMN separation variability. These factors might include differences across scanners in the multi-site ABCD study, higher levels of head motion in children, poorer grasp of task instructions in children compared to adults, and other such factors. A second possibility is that from a developmental perspective, critical aspects of the FPN/DMN network architecture supporting general intelligence are still immature in 9- to 10-year olds and there is still substantial neurodevelopmental change that lies ahead^96–100^. In this regard, it is noteworthy that functional connections involving FPN and DMN have been shown to be among most intensely maturing from early adolescence to young adulthood^79–81,101,102^. We will be able to clarify the relative roles of these two explanations in the coming years taking advantage of the longitudinal nature of the ABCD study, which will follow the baseline sample over the next ten years. Moreover, to the extent that the second explanation turns out to play an important role, we will be able to examine in detail the maturational changes in brain network architecture that facilitate the emergence of adult patterns of general intelligence functioning.

In sum, this study introduces separation between FPN and DMN as a novel brain network mechanism of general intelligence and a major locus of individual-differences. Our results invite systematic investigation into the neural, developmental, and genetic mechanisms of regulation of FPN/DMN separation in different cognitive tasks, across the lifespan, and in health and disease.

## Methods

### 1 HCP Task Analysis

#### 1.1 Data Acquisition and Preprocessing

We used data from the HCP-1200 release^51,52^ and all research was performed in accordance with relevant guidelines and regulations. Subjects provided informed consent, and recruitment procedures and informed consent forms, including consent to share de-identified data, were approved by the Washington University institutional review board. Subjects completed two runs each of seven scanner tasks across two fMRI sessions, using a 32-channel head coil on a 3T Siemens Skyra scanner (TR = 720ms, TE = 33.1ms, 72 slices, 2mm isotropic voxels, multiband acceleration factor = 8) with right-to-left and left-to-right phase encoding directions. Comprehensive details are available in the papers describing HCP’s overall neuroimaging approach^51,103^.

The main analysis was performed on an N-back working memory task, in which participants respond when the picture shown on the screen is the same as the one two trials back (=*2-back* condition) or the same as one shown at the start of the block (=*0-back* condition). Stimuli consisted of pictures of places, tools, faces, and body parts. Within each run, the 4 different stimulus types were presented in separate blocks. Also, within each run, ½ of the blocks use a 2-back working memory task and ½ use a 0-back working memory task. A 2.5 second cue indicates the task type (and target for 0-back) at the start of the block. Each of the two runs contains 8 task blocks (10 trials of 2.5 seconds each, for 25 seconds) and 4 fixation blocks (15 seconds). On each trial, the stimulus is presented for 2 seconds, followed by a 500 ms inter-task interval. 405 time points were collected per run, resulting in 4.86 minutes of data per run, or 9.72 minutes combining both runs.

An additional analysis was performed on the HCP Relational Task. In this task, participants identify the dimension along which a cue pair of objects differs and determine if a target pair differs along same dimension (=*relational* condition). Or they determine if a cue object matches a member of a target pair along a given dimension (=*match* condition). For both conditions, the subject responds yes or no using buttons. For the relational condition, the stimuli are presented for 3.5 seconds, with a 500 ms seconds ITI, and there are four trials per block. In the matching condition, stimuli are presented for 2.8 seconds, with a 400 ms ITI, and there are 5 trials per block. Each type of block (relational or matching) lasts a total of 18 seconds. In each of the two runs of this task, there are 3 relational blocks, 3 matching blocks and 3 16-second fixation blocks.

Data was preprocessed through the HCP minimally preprocessed pipeline, which is presented in detail by Glasser *et al.*^54^ and task details are described by Barch *et al*.^34^ Briefly, the pipeline includes gradient unwarping, motion correction, field-map distortion correction, brain-boundary based linear registration of functional to structural images, non-linear registration to MNI152 space, and grand-mean intensity normalization. Data then entered a surfaced-based preprocessing stream, followed by grayordinate-based processing, which involves data from the cortical ribbon being projected to surface space and combined with subcortical volumetric data. Left hemisphere surface, right hemisphere surface, and subcortical volume data from the grayordinate space were split and processed separately for all steps. Subject-level fixed-effects analyses were conducted using FEAT to estimate the average effects across runs within-participants.

#### 1.2 Inclusion/Exclusion Criteria

For the N-back analysis, subjects were eligible to be included if they had available MSMAll registered task data for both runs of the N-back task, full behavioral data for this task, no more than 25% of their volumes in each run exceeded a framewise displacement threshold of 0.5mm, and accuracy on both 0-back and 2-back conditions was at least 60%. These exclusions resulted in 920 subjects for the N-back fMRI analysis.

For the Relational analysis, subjects were eligible to be included if they had available MSMAll registered task data for both runs of the task, full behavioral data for this task, no more than 25% of their volumes in each run exceeded a framewise displacement threshold of 0.5mm, and accuracy on both the match and relational conditions was at least 40%. These exclusions resulted in 917 subjects for the Relational fMRI analysis.

#### 1.3 Constructing a General Intelligence Factor

**Figure 10:**
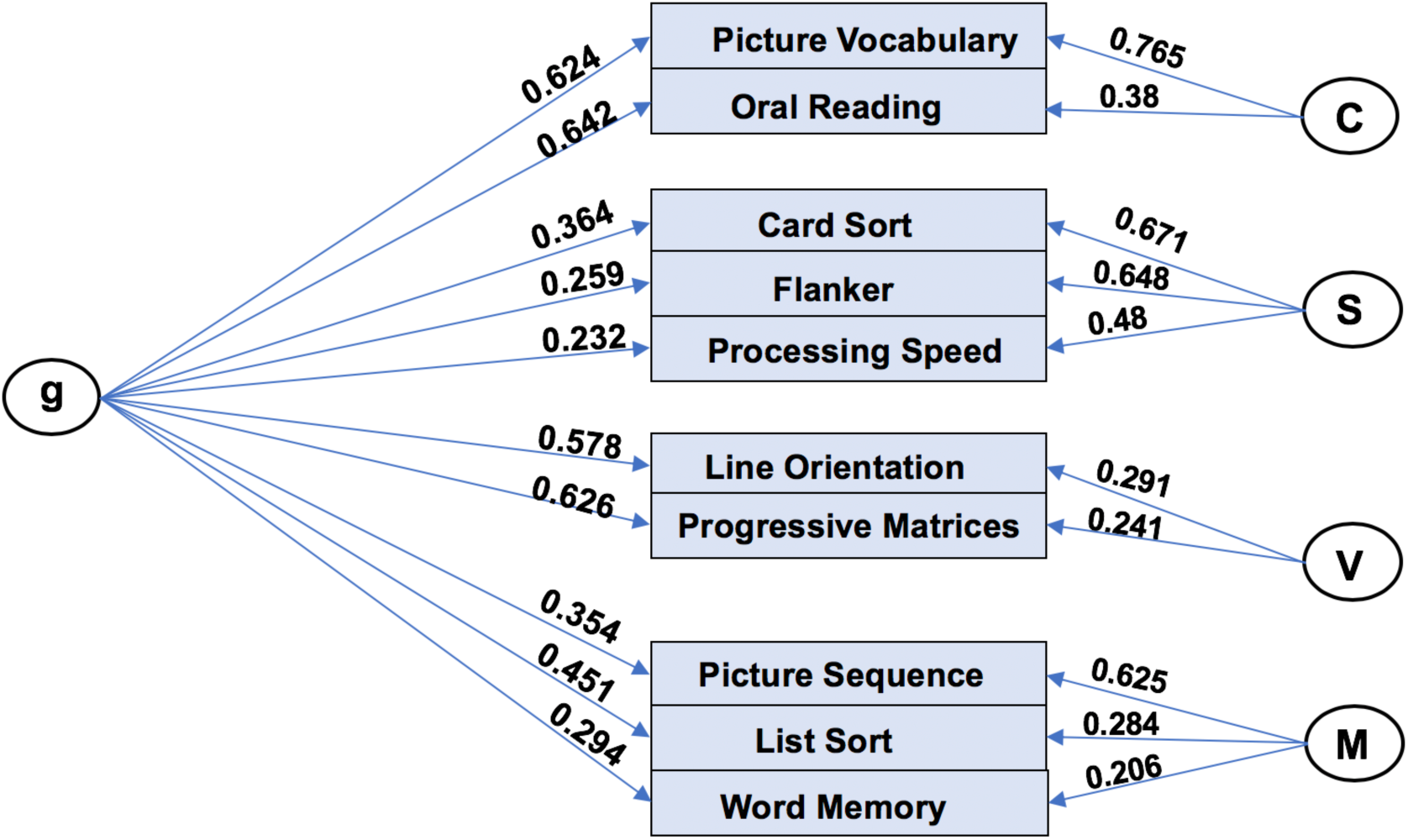
Bifactor Model Based on Ten Behavioral Tasks from the HCP Dataset with General Factor (“g”) and Four Group Factors. C=Crystallized Intelligence, S=Processing Speed, V=Visuospatial Ability, M=Memory.

We conducted an exploratory factor analysis utilizing the approachß and associated code made available by Dubois and colleagues (https://github.com/adolphslab/HCP_MRI-behavior), who recently investigated prediction of intelligence from resting state fMRI in the HCP dataset^58^. Unadjusted scores from ten cognitive tasks for 1181 HCP subjects were included in the analysis (subjects with missing data or MMSE < 26 were excluded), including seven tasks from the NIH Toolbox (Dimensional Change Cart Sort, Flanker Task, List Sort Test, Picture Sequence Test, Picture Vocabulary Test, Pattern Completion Test, Oral Reading Recognition Test) and three tasks from the Penn Neurocognitive Battery (Penn Progressive Matrices, Penn Word Memory Test, Variable Short Penn Line Orientation Test), with additional details supplied in ^58^.

We applied Dubois and colleagues’ code to this data, which uses the omega function in the psych (v 1.8.4) package^104^ in R (v3.4.4). In particular, the code performs maximum likelihood-estimated exploratory factor analysis (specifying a bifactor model), oblimin factor rotation, followed by a Schmid-Leiman transformation^105^ to find general factor loadings. The resulting bifactor model exhibited very good fit to the data (CFI=0.99; RMSEA=0.03; SRMR=0.02; BIC=- 0.52), which is substantially better than a single factor model (CFI=0.72; RMSEA=0.14; SRMR=0.09; BIC=591.2). The general factor accounted for 59% of the variance in task scores, group factors accounted for 18% of the variance, and 15% of the variance was unexplained (see Dubois and colleagues^58^ for additional discussion of the fit of this bifactor model).

To assess reliability of general intelligence factor scores, in a separate analysis, we re-ran the factor analysis excluding 46 subjects that had Test/Retest sessions available. We then estimated factor scores for both sessions for these subjects and calculated test/retest reliability via intraclass correlation (we used ICC(3,1) in the Shrout and Fleiss scheme^62^).

### 2 HCP Resting State Analysis

#### 2.1 Data Acquisition, Preprocessing, and Connectome Generation

Data used was from the HCP-1200 release^51,52^. Four runs of resting state fMRI data (14.4 minutes each; two runs per day over two days) were acquired using the same sequence described above in Methods, §1.1. Processed volumetric data from the HCP minimal preprocessing pipeline including ICA-FIX denoising were used. Full details of these steps can be found in Glasser^103^ and Salimi-Korshidi^106^.

Data then went through a number of resting state processing steps, including a motion artifact removal steps comparable to the type B (i.e., recommended) stream of Siegel et al.^107^. These steps include linear detrending, CompCor to extract and regress out the top 5 principal components of white matter and CSF^108^, bandpass filtering from 0.1-0.01Hz, and motion scrubbing of frames that exceed a framewise displacement of 0.5mm. We next calculated spatially-averaged time series for each of 264 4.24mm radius ROIs from the parcellation of Power et al.^109^. We then calculated Pearson’s correlation coefficients between each ROI. These were then were transformed using Fisher’s r to z-transformation.

#### 2.2 Inclusion/Exclusion Criteria

Subjects were eligible to be included if they had structural T1 data and had 4 complete resting state fMRI runs (14m 24s each). Subjects with more than 10% of frames censored were excluded from further analysis. Subjects had to have full necessary behavioral data and not be one of the subjects with retest data. This resulted in 910 subjects.

#### 2.3 Train/Test Split

We previously constructed a train/test split of the usable HCP resting state data, which is described in detail in our previous report^69^. In brief, this split included 810 train subjects and 100 test subjects unrelated to the train subjects and to each other. For the purposes of predicting task activation signatures from resting state for the present study, we intersected subjects with usable task data for each task with the resting state train/test split. For the prediction of FPN/DMN separation in the N-back task, this yielded 834 subjects total; 744 in the train set and 90 unrelated subjects in the test set. For the Relational task, this yielded 819 subjects total; 734 in the train set and 85 unrelated subjects in the test set.

#### 2.4 Brain Basis Set Modeling

To generate predictions of phenotypes from a basis set consisting of *k* components, we used Brain Basis Set (BBS) modeling, a predictive modeling approach validated in our previous studies^59,69,70^. This approach is similar to principal component regression^110,111^, with an added predictive modeling element. In a training partition, we calculate the expression scores for each of *k* components for each subject by projecting each subject’s connectivity matrix onto each component. We then fit a linear regression model with these expression scores as predictors and the phenotype of interest as the outcome, saving **B**, the *k × 1* vector of fitted coefficients, for later use. In a test partition, we again calculate the expression scores for each of the *k* components for each subject. Our predicted phenotype for each test subject is the dot product of **B** learned from the training partition with the vector of component expression scores for that subject. We set k at 75 based on our prior study^69^ that showed that larger values tend to result in overfitting and worse performance.

#### 2.5 Consensus Component Maps for Visualization

To help convey overall patterns across the entire BBS predictive model, we constructed “consensus” component maps. We first fit a BBS model to the entire dataset consisting of all participants. We then multiplied each component map with its associated beta from this fitted BBS model. Next, we summed across all components yielding a single map, and thresholded the entries at *z*=2. The resulting map indicates the extent to which each connection is positively (red) or negatively (blue) related to the outcome variable of interest.

### 3 ABCD Task Analysis

#### 3.1 Data Acquisition and Preprocessing

The ABCD study is a multisite longitudinal study established to investigate how individual, family, and broader socio-cultural factors shape brain development and health outcomes. The study has recruited 11,875 children between 9-10 years of age from 21 sites across the United States for longitudinal assessment. The study conforms to the rules and procedures of each site’s Institutional Review Board, and all participants provide informed consent (parents) or informed assent (children). At each assessment wave, children undergo assessments of neurocognition, physical health, and mental health, and also participate in structural and functional neuroimaging. Detailed description of recruitment procedures^112^, assessments^113^, and imaging protocols^114^ are available elsewhere. The ABCD data repository grows and changes over time. The ABCD data used in this report came from NDA Study 576, DOI 10.15154/1412097, which can be found at https://ndar.nih.gov/study.html?id=576.

Imaging protocols were harmonized across ABCD sites and scanners. For the current analysis, minimally preprocessed fMRI data from the curated ABCD annual release 1.1 were used, and full details are described in ^115^. This data reflects the application of the following steps: i) gradient-nonlinearity distortions and inhomogeneity correction for structural data; and ii) gradient-nonlinearity distortion correction, rigid realignment to adjust for motion, and field map correction for functional data. Additional processing steps were applied by our group using SPM12, including co-registration, segmentation and normalization using the CAT12 toolbox and DARTEL, smoothing with a 6mm FWHM Gaussian kernel, and application of ICA-AROMA^116^.

The ABCD N-back task is broadly similar to the HCP N-back task in having high and low memory load conditions (2 back and 0 back). The main difference is that the stimuli in the ABCD N-back task consist of faces (happy, fearful and neutral) and places, but no tools and body parts. The task includes two runs of eight blocks each (4.93 minutes per run, 9.87 minutes across both runs). Each block consists of 10 trials (2.5 s each) and 4 fixation blocks (15 s each). Each trial consists of a stimulus presented for 2 s, followed immediately by a 500 ms fixation cross, with 160 trials total.

#### 3.2 Inclusion/Exclusion Criteria

Subjects were eligible to be included if they had available task data for at least one run of the N-back task, full necessary behavioral data, at least 3 minutes of data in each run was below a framewise displacement threshold of 0.5mm, and accuracy on both 0-back and 2-back conditions was at least 60%. Additionally, visual inspections were performed of the coregistration and normalization steps and subjects with poor coregistration and/or normalization were excluded. The auto-generated first level masks from SPM were compared to a canonical brain mask in MNI space to exclude subjects with excessive signal loss, and subjects with less than 90% coverage of the canonical mask were excluded. Finally, we excluded those sites that had fewer than 75 subjects that passed the above criteria in order to allow sufficient subjects at each site for regression-based removal of site effects. These exclusions resulted in 1240 subjects from 10 sites for the task fMRI analysis. To account for site effects in the data, we regressed the linear effect of site voxel-wise from included subjects’ N-back task contrast maps.

#### 3.3 Constructing a General Intelligence Factor

**Figure 11:**
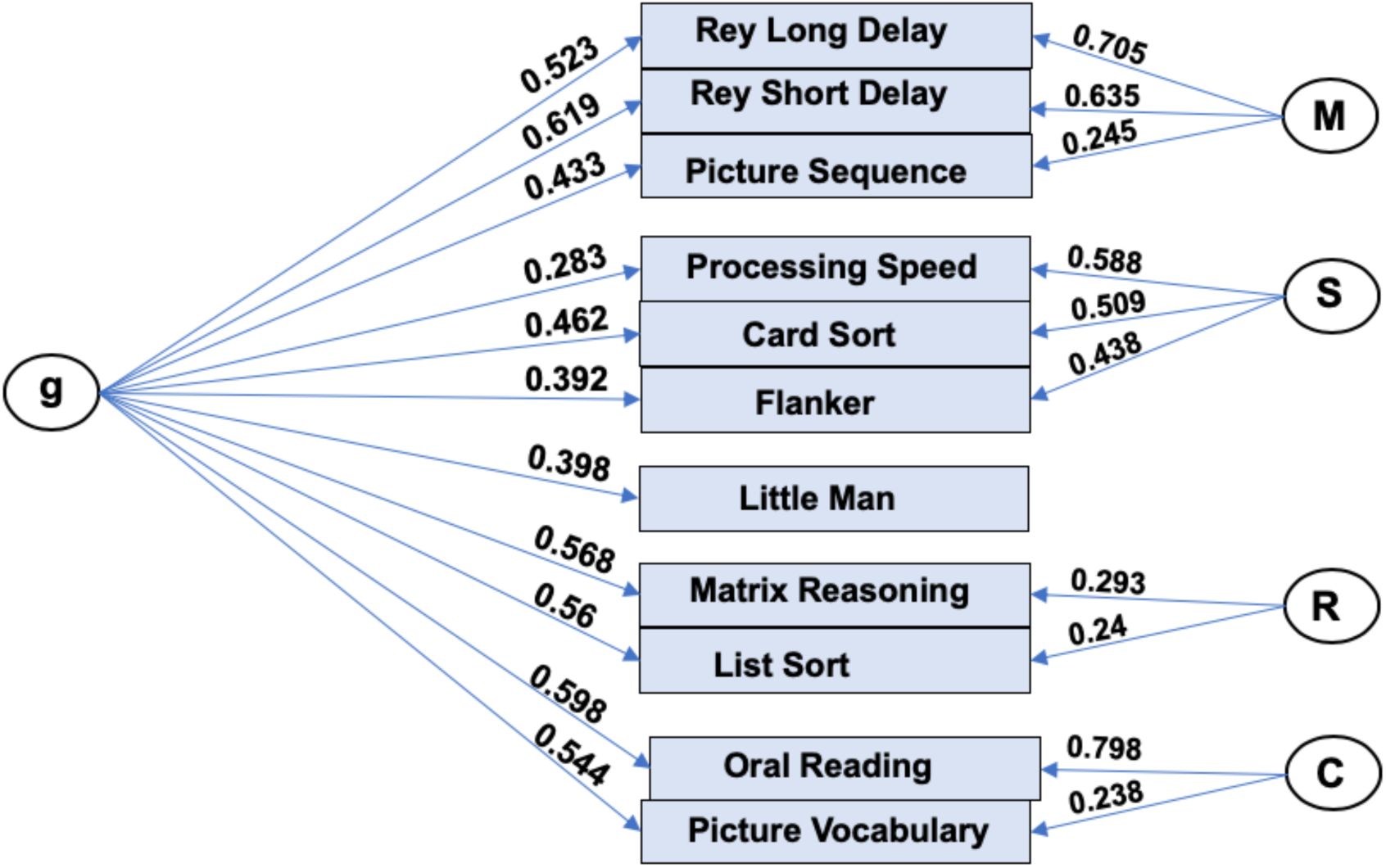
Bifactor Model Based on Eleven Behavioral Tasks from the ABCD Dataset with General Factor (“g”) and Four Group Factors. C=Crystallized Intelligence, S=Processing Speed, R=Reasoning, M=Memory.

As above in the HCP data, we applied Dubois and colleagues’ exploratory factor analysis code to behavioral task data to find a general intelligence factor. For the ABCD dataset, the variables included in the analysis were unadjusted scores from the NIH toolbox (Dimensional Change Card Sort, Flanker Task, List Sort Test, Picture Sequence Test, Picture Vocabulary Test, Pattern Completion Test, Oral Reading Recognition Test) as well as total correct from the Short Delay and Long Delay of the Rey Auditory Verbal Leaning Task, WISC-V Matrix Reasoning total raw score, and number correct on the Little Man Task. The bifactor model exhibited very good fit to the data (CFI=0.99; RMSEA=0.03; SRMR=0.01; BIC=2.3), which is substantially better than a single factor model (CFI=0.74; RMSEA=0.14; SRMR=0.07; BIC=2906.7). The general factor accounted for 66% of the variance in task scores, group factors accounted for 16% of the variance, and 13% of the variance was unexplained.

### 4 ABCD Resting State Analysis

#### 4.1 Data Acquisition, Preprocessing, and Connectome Generation

Data acquisition/preprocessing for the resting state data follows the steps described for task data above in section 3.1, which includes application of ICA-AROMA. Resting state fMRI was acquired in four separate runs (5 minutes per run, 20 minutes total). Resting state-specific processing steps applied including linear detrending, CompCor to regress the top 5 components of CSF and WM signal^108^, bandpass filtering from 0.1-0.01Hz, and motion scrubbing of frames that exceed a framewise displacement of 0.5mm. We next applied the parcellation of Power et al.^117^, calculated Pearson’s correlation coefficients between each ROI, and applied Fisher’s *r* to *z*-transformation.

#### 4.2 Inclusion/Exclusion

There were 4521 subjects in the ABCD Release 1.1 dataset. Of these, 3575 subjects had usable T1w images and one or more resting state runs that passed ABCD quality checking standards (fsqc_qc = 1). Next, 3544 passed preprocessing and were subsequently visually checked for registration and normalization quality, where 197 were excluded for poor quality. Motion was assessed based on number of frames censored, with a framewise displacement threshold of 0.5mm, and only subjects with two or more runs with at least 4 minutes of good data were included (*n*=2757). To remove unwanted sources of dependence in the dataset, only one sibling was randomly chosen to be retained for any family with more than 1 sibling (*n*=2494). Finally, in order to implement leave-one-site-out cross validation, sites with fewer than 75 subjects that passed these quality checks were dropped, leaving 2206 subjects across 15 sites to enter the PCA step of BBS predictive modeling. For the purposes of predicting task activation signatures from resting state, this dataset was then intersected with the included ABCD task dataset above to yield 946 subjects from 10 sites.

#### 4.3 Brain Basis Set Modeling

The Brain Basis Set modeling used for the ABCD data followed the same methodology as reported for the HCP dataset above, with the exception that in ABCD we utilized a leave-one-site-out cross validation strategy instead of a single, independent training/testing split. The BBS procedure was repeated 10 times, each time holding one site out of the training data to be used as test data. Correlations were calculated between observed and predicted phenotype scores for each fold, transformed with a Fisher’s R to Z transformation, averaged across folds, and then transformed back to correlations with a Fisher’s Z to R.

#### 4.4 Permutation Testing Framework

To assess the statistical significance of brain basis set models (BBS) in the ABCD dataset, we used non-parametric permutation methods. The distribution under chance of correlations between BBS-based predictions of FPN/DMN separation values and observed FPN/DMN separation values was generated by randomly permuting the subjects’ FPN/DMN separation values 10,000 times. At each iteration, we performed the leave-one-site out cross validation procedure described above, which includes refitting BBS models at each fold of the cross-validation. We then recalculated the average correlation across folds between predicted versus actual neurocognitive scores. The average correlation across folds that was actually observed was located in this null distribution in terms of rank, and statistical significance was set as this rank value divided by 10,000.

Since the BBS models fit at each iteration of the permutation test included covariates (mean FD and mean FD squared), the procedure of Freedman and Lane was followed^118^. In brief, a BBS model was first estimated with nuisance covariates alone, residuals were formed and were permuted. The covariate effect of interest was then included in the subsequent model, creating an approximate realization of data under the null hypothesis, and the statistical test of interest was calculated on this data (see FSL Randomise http://fsl.fmrib.ox.ac.uk/fsl/fslwiki/Randomise/Theory for a neuroimaging implementation).

## Acknowledgments

HCP data were provided [in part] by the Human Connectome Project, WU-Minn Consortium (Principal Investigators: David Van Essen and Kamil Ugurbil; 1U54MH091657) funded by the 16 NIH Institutes and Centers that support the NIH Blueprint for Neuroscience Research; and by the McDonnell Center for Systems Neuroscience at Washington University.

ABCD data used in the preparation of this article were obtained from the Adolescent Brain Cognitive Development (ABCD) Study (https://abcdstudy.org), held in the NIMH Data Archive (NDA). This is a multisite, longitudinal study designed to recruit more than 10,000 children age 9-10 and follow them over 10 years into early adulthood. The ABCD Study is supported by the National Institutes of Health and additional federal partners under award numbers U01DA041022, U01DA041028, U01DA041048, U01DA041089, U01DA041106, U01DA041117, U01DA041120, U01DA041134, U01DA041148, U01DA041156, U01DA041174, U24DA041123, U24DA041147, U01DA041093, and U01DA041025. A full list of supporters is available at https://abcdstudy.org/federal-partners.html. A listing of participating sites and a complete listing of the study investigators can be found at https://abcdstudy.org/Consortium_Members.pdf. ABCD consortium investigators designed and implemented the study and/or provided data but did not necessarily participate in analysis or writing of this report. This manuscript reflects the views of the authors and may not reflect the opinions or views of the NIH or ABCD consortium investigators.

This work was supported by the following grants from the United States National Institutes of Health, the National Institute on Drug Abuse, and the National Institute on Alcohol Abuse and Alcoholism: R01MH107741 (CS), U01DA041106 (CS), In addition, CS was supported by a grant from the Dana Foundation David Mahoney Neuroimaging Program. This research was supported in part through computational resources and services provided by Advanced Research Computing at the University of Michigan, Ann Arbor.

## Supplement

### Supplemental Results

**Figure S1:**
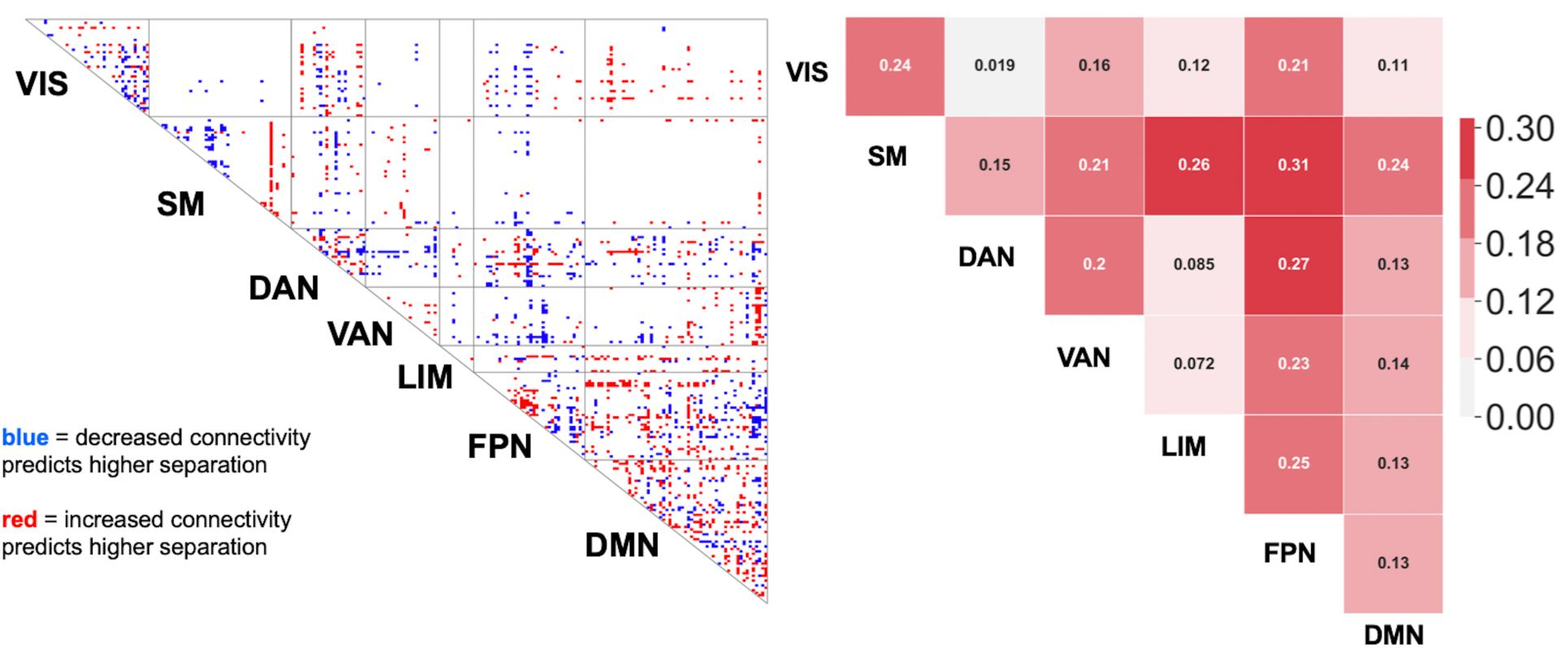
Connections During Resting State That Are Predictive of FPN/DMN Separation During the Relational Task in the HCP sample. We found a predictive model trained on resting state data predicted change in FPN/DMN separation between the match and relational conditions (“delta FPN/DMN separation”) with a correlation of 0.21, which is statistically significant (p = 0.05). (Left Panel) Consensus connectome showing connections weighted highly in the predictive model. Similar to the HCP dataset, the consensus connectome showed a preponderance of connections involving FPN and DMN (FPN and DMN make up 12.9% percent of connections in the connectome but constitute 39.5% of the suprathreshold connections). (Right Panel) Table showing predictive success in follow-up analyses that dropped all networks except two. Several network pairs involving FPN and DMN showed strong predictive success, with somatomotor network also playing an unexpectedly important role.

**Figure S2:**
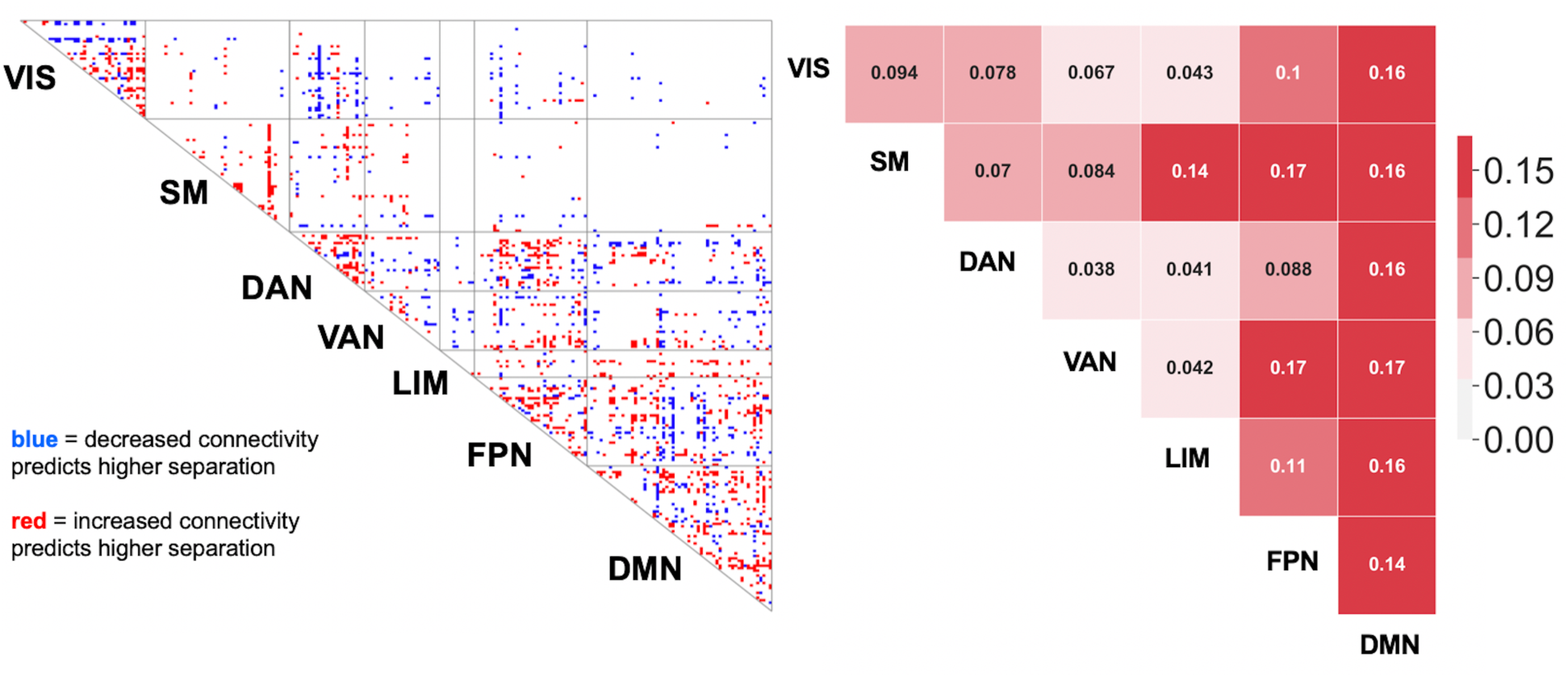
Connections During Resting State That Are Predictive of FPN/DMN Separation During the N-Back Task in the ABCD Sample. Using leave-one-site-out cross-validation, we found a predictive model trained on resting state data predicted change in FPN/DMN separation between the 0-back and 2-back conditions (“delta FPN/DMN separation”) with a correlation of 0.17, which is statistically significant (permutation p value = 0.0004). (Left Panel) Consensus connectome showing connections weighted highly in the predictive model. Similar to the HCP dataset, the consensus connectome showed a preponderance of connections involving FPN and DMN (FPN and DMN make up 12.9% percent of connections in the connectome but constitute 40.0% of the suprathreshold connections). (Right Panel) Table showing predictive success in follow-up analyses that dropped all networks except two. Several network pairs involving FPN and DMN showed strong predictive success.

